# Presaccadic attention depends on eye movement direction and is related to V1 cortical magnification

**DOI:** 10.1101/2022.12.15.520489

**Authors:** Nina M. Hanning, Marc M. Himmelberg, Marisa Carrasco

## Abstract

With every saccadic eye movement, humans bring new information into their fovea to be processed with high visual acuity. Notably, perception is enhanced already before a relevant item is foveated: During saccade preparation, *presaccadic attention* shifts to the upcoming fixation location, which can be measured via behavioral correlates such as enhanced visual performance or modulations of sensory feature tuning. The coupling between saccadic eye movements and attention is assumed to be robust and mandatory, and considered a mechanism facilitating the integration of pre- and post-saccadic information. However, until recently it had not been investigated as a function of saccade direction. Here, we measured contrast response functions during fixation and saccade preparation in male and female observers and found that the pronounced response gain benefit typically elicited by presaccadic attention is selectively lacking before upward saccades at the group level – some observers even showed a cost. Individual observers’ sensitivity before upward saccades was negatively related to their amount of surface area in primary visual cortex representing the saccade target, suggesting a potential compensatory mechanism that optimizes the use of the limited neural resources processing the upper vertical meridian. Our results raise the question how perceptual continuity is achieved and upward saccades can be accurately targeted despite the lack of – theoretically required– presaccadic attention.

**Significance Statement:** When we make a saccadic eye movement to a target location in the visual field, perception improves at the saccade target, already before the eyes start moving. This benefit afforded by presaccadic attention is thought to be mandatory and independent of eye movement direction. We show that this is not the case; moving our eyes horizontally or downwards, but not upwards, enhances contrast sensitivity. At the neural level, however, humans with less V1 cortical tissue representing the target location for upwards saccades have some presaccadic enhancement. The finding that presaccadic attention is dependent upon eye movement direction challenges the view that the presaccadic benefit is automatic and mandatory in nature.

## Introduction

As the high acuity of the fovea markedly decreases with eccentricity, we constantly make saccadic eye movements to actively explore the visual world and gather information. To maintain a continuous percept across saccades, the human visual system must seamlessly integrate blurry peripheral information with its high-acuity equivalent, once brought into the fovea by a saccade. To achieve a smooth transition, we preferentially process visual information at the future eye fixation already before the eyes start moving: During saccade preparation, ‘presaccadic attention’ is deployed to the saccade target, where it for example enhances visual sensitivity (Kowler et al., 1995; Deubel and Schneider, 1996; Montagnini and Castet, 2007; Rolfs and Carrasco, 2012) and sharpens orientation and spatial frequency tuning (Li et al., 2016, 2019; Ohl et al., 2017; Kroell and Rolfs, 2021). These presaccadic perceptual modulations render the peripheral information at the future eye fixation more fovea-like, easing the integration of pre-and post-saccadic visual input in support of perceptual continuity across saccades. Presaccadic attention is considered robust and mandatory (Deubel and Schneider, 1996; Li et al., 2019; Kreyenmeier et al., 2020; Hanning et al., 2022b); sensitivity at non-target locations –including the currently still foveated location (Hanning and Deubel, 2022)– inevitably decreases just before saccade onset (Montagnini and Castet, 2007; Deubel, 2008; Collins et al., 2010).

The assumed neural basis of presaccadic perceptual modulations are feedback signals from oculomotor structures (e.g., frontal eye fields, FEF; superior colliculus, SC) to early visual cortices (Ekstrom et al., 2008; Bisley and Mirpour, 2019). Subthreshold micro-stimulation of FEF or SC (which elicits a saccade when stimulated above threshold) changes activity in visual cortex (Moore and Armstrong, 2003) and enhances sensitivity at the movement field of the stimulated neurons (Moore and Fallah, 2004; Müller et al., 2005), resembling the behavioral correlates of presaccadic attention.

The strong functional coupling of oculomotor programming and attention is undisputed and neural and behavioral effects of presaccadic attention are thought to be ubiquitous throughout the visual field – independent of where the saccade is executed. Visual performance, however, is heterogeneous around the visual field (Himmelberg et al., 2023b): At a fixed eccentricity, it is better along the horizontal than vertical meridian, and along the lower than upper vertical meridian. These polar angle asymmetries emerge for various dimensions, e.g., contrast sensitivity, spatial resolution (Abrams et al., 2012; Barbot et al., 2021), and are robust enough to not be attenuated by covert attention (e.g., Carrasco et al., 2001; Roberts et al., 2016; Purokayastha et al., 2021), deployed without concurrent eye movements. Presaccadic attention even exacerbates them: measured at perceptual threshold, saccade preparation enhances contrast sensitivity before horizontal and downward, but not upward saccades (Hanning et al., 2022a) (**Figure 1a**).

**Figure 1.**
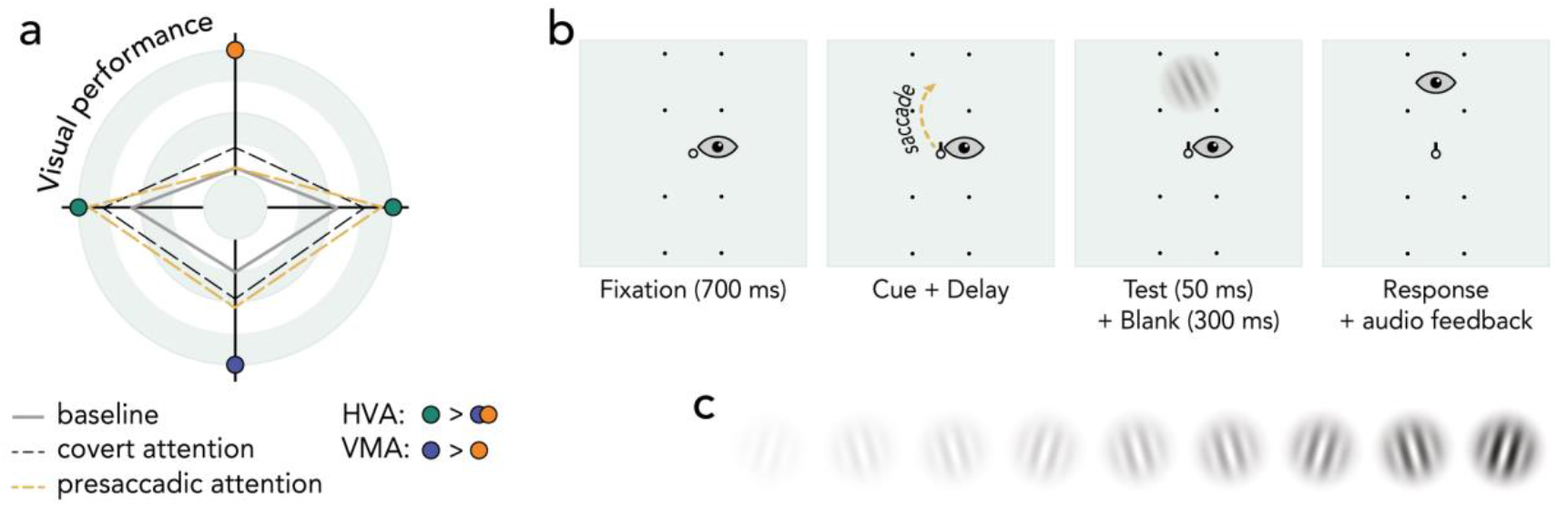
Background and experimental design. (**a**) Schematic representation of polar angle performance asymmetries. Points further from the center represent higher performance. Baseline visual performance (during fixation) is better along the horizontal than vertical meridian (HVA), and along the lower than upper vertical meridian (VMA). Covert spatial (Carrasco et al., 2001; Cameron et al., 2002; Talgar and Carrasco, 2002; Roberts et al., 2016; Purokayastha et al., 2021) and temporal attention (Fernández et al., 2019), tested under similar stimulus conditions, benefit performance to the same degree around the visual field. Presaccadic attention, however, exacerbates the asymmetries by enhancing contrast sensitivity at all locations except the upper vertical meridian(Hanning et al., 2022a). (**b**) Experimental task. Vertical *saccade* condition. After a fixation period, a central direction cue (black line) indicated the saccade target 8° above or below central fixation, marked by black dots (note that in the horizontal conditions stimuli were placed 8° left and right). ~100ms after direction cue onset (delay adjusted for each observer), a test Gabor was presented at the saccade target; observers reported its orientation after saccade offset. In the *fixation* condition, two lines cued both placeholder locations; observers were instructed to maintain fixation. See **Movie 1** for a demonstration of the trials sequence. (**c**) Visualization of the different contrast levels (5-95%, log-spaced) tested to measure contrast response functions.

This finding is surprising: (I) Presaccadic attention is assumed to be mandatorily deployed before any saccade being made (Shepherd et al., 1986; Deubel and Schneider, 1996; Hanning et al., 2022b); (II) By rendering peripheral representations at the saccade target more fovea-like, presaccadic attention enables a smooth integration of pre- and post-saccadic information and thus aids perceptual continuity (Herwig, 2015; Rolfs, 2015; Kwon et al., 2019)–a mechanism that should be indispensable also for upward saccades; (III) Sensitivity at the upper vertical meridian (during fixation) is typically lowest (review: Himmelberg et al., 2023b), thus perceptual performance could improve there the most.

Our former study measured presaccadic attention at perceptual threshold (Hanning et al., 2022a), but presaccadic attention primarily modulates performance at high stimulus contrasts, where performance asymptotes (Li et al., 2021b; Hanning et al., 2023). To optimally investigate the extent of presaccadic attention as a function of saccade direction, here we measured full contrast response functions and evaluated presaccadic response gain around the visual field. Additionally, we obtained fMRI-derived retinotopic maps for our observers from the NYU Retinotopy Dataset (Himmelberg et al., 2021). Across individuals, greater local V1 surface area encoding a visual field location benefits perception at that location (Duncan and Boynton, 2003; Himmelberg et al., 2022). We explored whether individual differences in the amount of local V1 surface area encoding the saccade target region are related to the inconsistent magnitude of individual observers’ presaccadic enhancement, which at the group level was lacking before upward saccades.

## Materials & Methods

In this study, we compared 2AFC orientation discrimination performance during fixation –a neutral cue instructed observers to maintain fixation throughout the trial– and saccade preparation –observers made an immediate saccade to a centrally cued target location 8° left, right, above, or below fixation (**Figure 1b & Supplemental Movie 1**; *Experimental design*). We varied the contrast of the test grating on a trial-by-trial basis across the full contrast range (**Figure 1c**), to measure presaccadic modulations of the contrast response function (CRF) at the saccade target. Observers’ gaze positions were monitored continuously. Importantly, test presentation time was adjusted separately to each individual observer’s horizontal and vertical saccade latencies (*Stimulus timing*), to ensure a measurement right before saccade onset, where presaccadic attention has its maximum effect (Deubel, 2008; Rolfs and Carrasco, 2012; Li et al., 2016; Hanning et al., 2018, 2019a). Only trials in which the test was presented during the last 100ms before saccade onset were included in the analysis. We adjusted the tilt angle (clockwise or counter-clockwise relative to vertical) in a pre-test (*Tilt titration*) to compensate for polar angle asymmetries and ensure equal task difficulty around the visual field during fixation – and thus equal room for a presaccadic benefit.

### Observers

Eight observers (4 female, 4 male; aged 21–33 years, including two authors: NMH and MMH) with normal or corrected-to-normal vision participated in the study. We chose a sample size in the typical range of studies investigating presaccadic attention (Montagnini and Castet, 2007; Ohl et al., 2017; Hanning et al., 2019b; Li et al., 2019; Kreyenmeier et al., 2020; Li et al., 2021b; Hanning and Deubel, 2022). All observers provided written informed consent, and (except for the two authors) were naive to the purpose of the experiment. The protocols for the study were approved by the University Committee on Activities involving Human Subjects at New York University and all experimental procedures were in accordance with the Declaration of Helsinki.

### Setup

Observers sat in a dimly illuminated room with their head stabilized by a chin and forehead rest and viewed the stimuli at 57cm distance on a gamma-linearized 20-inch ViewSonic G220fb CRT screen (Brea, CA, USA) with a spatial resolution of 1,280 by 960 pixels and a vertical refresh rate of 85 Hz. Gaze position of the dominant eye was recorded using an EyeLink 1000 Desktop Mount eye tracker (SR Research, Osgoode, Ontario, Canada) at a sampling rate of 1 kHz. Manual responses were recorded via a standard keyboard. An Apple iMac Intel Core 2 Duo computer (Cupertino, CA, USA) running Matlab (MathWorks, Natick, MA, USA) with Psychophysics (Brainard, 1997; Pelli, 1997) and EyeLink (Cornelissen et al., 2002) toolboxes, controlled stimulus presentation and response collection.

### Experimental design

The experiment comprised two experimental conditions –*saccade* and *fixation*– tested at the two meridians–horizontal and vertical– in separate experimental blocks. Observers fixated a central fixation target (black ring; ~0 cd/m^2^, diameter 0.35°, width 0.1° of visual angle) on gray (~26 cd/m^2^) background (see **Figure 1b**). Two placeholders indicated the isoeccentric locations of the upcoming stimuli (and potential saccade targets), either 8° left and right, or above and below fixation (depending on the tested meridian). Each placeholder comprised four dark gray dots (diameter 0.1°), forming the corners of a square (diameter 4.5°). The trial started once we detected stable fixation within a 1.75° radius virtual circle centered on the fixation target.

In *saccade* blocks, after a 700ms fixation period, a central direction cue (black line; ~0 cd/m^2^, length 0.175°, width 0.1°) pointed to one of the opposing placeholders (randomly selected), cueing the saccade target. Observers were instructed to look as fast and precisely as possible to the center of the indicated placeholder. Approximately 100ms after cue onset (inter-stimulus interval adjusted based on each observer’s direction specific movement latency, see *Stimulus timing*), a Gabor grating (4cpd, random phase, Gaussian envelope s = 0.5°), which was slightly tilted relative to vertical (see *Tilt titration*), appeared for ~35ms at the cued saccade target. Gabor contrast was varied on a trial-by-trial basis following the methods of constant stimuli (Michelson contrast, 9 log-spaced steps from 5% to 95%, see **Figure 1c**). Note that the grating was presented within the movement latency, i.e., when gaze still rested at fixation. 400ms after stimuli offset (well after the eye had landed at the saccade target), we played a high- or low-pitch response sound to specify the location that had contained the Gabor patch. Observers indicated their orientation judgement via button press (clockwise or counterclockwise, two-alternative forced choice) and were informed that the orientation report was non-speeded. They received auditory feedback for incorrect responses. Stimulus parameters and timing for the *fixation* blocks were identical to the *saccade* blocks, with one difference: two (rather than one) black direction cue lines appeared, pointing to both placeholder locations and observers were instructed to keep eye fixation while their gaze was monitored.

Observers performed multiple sessions of 12 experimental blocks each: 4 horizontal and 8 vertical blocks (counterbalanced within sessions), of which half were *fixation* and half were *saccade* blocks, randomly interleaved. Note that less experimental blocks were dedicated to testing the horizontal than vertical meridian because we collapsed data across the left and right test location (but separately analyzed upward and downward locations). Each block comprised 105 trials. We monitored gaze position online and controlled for correct eye fixation, i.e. gaze remaining within 1.75° from the central fixation target until (a) response cue onset (*fixation* blocks) or (b) direction cue onset (*saccade* blocks). Trials in which gaze deviated from fixation were aborted and repeated at the end of each block. In *saccade* blocks we also repeated trials with too short (<100ms) or long (>400ms) saccade latency, or incorrect eye movements (initial saccade landing beyond 2.5° from the indicated target). Overall data collection for an observer was stopped once at least 50 trials per stimulus contrast level and experimental condition were acquired after offline trial exclusion (see *Eye data preprocessing*).

### Tilt titration

To match overall task difficulty to each observers’ visual sensitivity and to account for sensitivity differences around the visual field, we titrated the tilt angle (± 0.125° – 5° relative to vertical) separately for each observer and test position (horizontal, upper, lower) with best PEST (Pentland, 2010) (using custom code [https://github.com/michaeljigo/palamedes_wrapper] that ran subroutines implemented in the Palamedes toolbox (Prins and Kingdom, 2018)). Using the procedure of the *fixation* condition, we concurrently ran 3 independent adaptive procedures (each comprising 36 trials) for each test location, to determine the tilt angle at which observers ‘orientation discrimination performance at the highest contrast level (95%) was ~d′ = 2. The location specific tilt angle used for the first main experimental session (same for *saccade* and *fixation* blocks) was computed by averaging medians of the last 5 trials of each individual staircase across the 3 staircases per location. Following each experimental session, the tilt angle for each location was adjusted as needed to ensure comparable accuracy across locations in the fixation condition. Note that the tilt angle for a given test location was the same for fixation and saccade blocks within an experimental session. Tilt angles did not correlate with the observed presaccadic benefit for any tested saccade direction (horizontal: *r*(6)=-.48, *p*=0.224; downward: *r*(6)=.65, *p*=0.080; upward: *r*(6)=-.24, *p*=0.573).

### Eye data preprocessing

We scanned the recorded eye-position data offline and detected saccades based on their velocity distribution (Engbert and Mergenthaler, 2006) using a moving average over 20 subsequent eye position samples. Saccade onset and offset were detected when the velocity exceeded or fell below the median of the moving average by 3 standard deviations for at least 20ms. We included trials in which no blink occurred during the trial and correct eye fixation was maintained within a 1.75° radius centered on central fixation throughout the trial (*fixation* trials) or until cue onset (*saccade* trials). Moreover, we only included those eye movement trials in which the initial saccade landed within 2.5 from the required target location and in which the test signal was presented within 100ms before saccade onset (i.e., the saccade started only after test signal presentation, but not later than 100ms after signal offset). In total, we included 30,168 trials in the analysis of the behavioral results (on average 3,771 ± 114 (mean ± 1 SEM) trials per observer).

### Stimulus timing

The effect of presaccadic attention increases throughout saccade preparation and peaks shortly before saccade onset(Deubel, 2008; Rolfs and Carrasco, 2012; Li et al., 2016; Hanning et al., 2018, 2019a). Saccade latencies, however, vary as a function of saccade direction (Honda and Findlay, 1992; Tzelepi et al., 2010; Hanning et al., 2022a; Kwak et al., 2023; Liu et al., 2023), e.g., downward saccades are typically initiated ~25ms later than horizontal or upward saccades (also see **Figure 3d**). To measure and compare the effects of presaccadic attention equally close to saccade onset for all saccade directions, we matched the test presentation time relative to the saccade by adjusting the delay between saccade cue and test onset to each observer’s direction specific saccade latency. For the initial experimental session, we randomly selected the stimulus onset asynchrony (SOA) between 141ms and 235ms on a trial-by-trial basis. After each session, we evaluated vertical and horizontal saccade latencies and adjusted the SOA-range to only include those SOAs for which test presentation fell in the desired presaccadic window (test offset -100 to 0ms relative to saccade onset) in at least 70% of trials. This adjusted SOA-range was used for both fixation and saccade blocks in the upcoming session.

### Quantification & Statistical analysis

Orientation discrimination performance, indexed by visual sensitivity (d-prime; d’ = *z*(hit rate) - *z*(false alarm rate)), was measured as a function of stimulus contrast using the method of constant stimuli (9 log-spaced Michelson contrast levels: 5, 10, 14, 20, 27, 37, 51, 70, 95%). We arbitrarily defined counter-clockwise responses to counter-clockwise oriented gratings as hits and counter-clockwise responses to clockwise oriented gratings as false-alarms (Huihui et al., 2019; Jigo and Carrasco, 2020; Li et al., 2021b). To avoid infinite values when computing d’, we substituted hit and false alarm rates of 0 and 1 by 0.01 and 0.99, respectively (Wollenberg et al., 2018; Hanning et al., 2019a, 2019b; Hanning and Deubel, 2020).

As only *saccade* trials with test offset occurring within the last 100ms prior to saccade onset were included in the sensitivity analysis (see *Eye data preprocessing*), trials with certain SOAs (see *Stimulus timing*) were more likely to contribute to the averaged visual sensitivity per contrast level. To ensure identical timing parameters in the fixation condition (in which no trials were excluded based on temporal criteria), for each observer we first calculated d’ for each stimulus contrast level separately for each used SOA, before computing a weighted average per contrast level, that matched the respective contrast level’s SOA distribution at the respective test location of the saccade condition.

To evaluate asymptotic contrast sensitivity, we obtained contrast response functions for each condition and test location by fitting each observer’s data with Naka-Rushton functions(Naka and Rushton, 1966), parameterized as d′(C) = d_max_C^n^ / (C^n^ + C^n^), where C_50_ is the contrast level, d_max_ is the asymptotic performance, C_50_ is the semi-saturation constant (contrast level corresponding to half the asymptotic performance), and n determines the slope of the function. The error was minimized using a least-squared criterion; d_max_ and c_50_ were free parameters, n was fixed. Contrast levels were log-transformed prior to fitting. A change in d_max_ indicates a response gain change and a change of C_50_ indicates a contrast gain change. We used repeated measures ANOVAs to assess statistical significance, followed by paired t-tests for post-hoc comparisons. All post-hoc comparisons were Bonferroni-corrected for multiple comparisons. All p-values for repeated-measures ANOVAs in which the assumption of sphericity was not met were Greenhouse-Geisser corrected. Parameter estimates for psychometric functions and all statistical tests were computed in MATLAB.

### fMRI Retinotopy & V1 Surface computation

Each observer’s population receptive field (pRF) data (Dumoulin and Wandell, 2008) and anatomical data were taken from the publicly-available NYU Retinotopy Dataset (Himmelberg et al., 2021). The pRF stimulus, MRI and fMRI acquisition parameters, MRI and fMRI preprocessing, the implementation of the pRF model, as well as the computation of V1 surface area representing wedge-ROIs along the cardinal meridians of the visual field are identical to those described in our previous work (Himmelberg et al., 2021, 2022). In brief, to calculate the V1 surface area representing each saccade target region, we defined wedge-ROIs in the visual field that were centered along each of the cardinal meridians. Each wedge-ROI was ±25° in width and extended 4°–12° of eccentricity (i.e., surrounding the saccade target and stimulus locations at 8° eccentricity, as including more data within the wedge-ROI leads to a more accurate estimate of V1 surface area). We calculated the amount of V1 surface area (in mm^2^) encapsulated by each of these wedge-ROIs by first generating 9 sub-wedge-ROIs (that are later combined to form the full wedge-ROI). Each sub-wedge-ROI was constrained to a narrow eccentricity band (9 log-spaced bands from 4°–12°). For each sub-wedge-ROI, the 25° angular border on the cortex was calculated by identifying the average distance of a selected pool of vertices whose pRF polar angle coordinates lied around the 25° polar angle pRF centers from each respective meridian. All vertices between a meridian and the 25° boundary were included in the sub-wedge-ROI. The sub-wedge-ROI was then used as a mask and overlaid on the anatomical surface, for which each vertex has an assigned surface area value (in mm^2^). The surface areas of the vertices within the sub-wedge-ROI mask were then summed to give a measurement of surface area for the respective sub-wedge-ROI. This was repeated for each sub-wedge ROI. Finally, the surface area of each sub-wedge-ROI was summed to compute the amount of surface area representing the full wedge-ROI representing a saccade target wedge in the visual field. This was repeated for each cardinal saccade target wedge. For the current study, the surface areas of the horizontal (left and right) saccade target wedge-ROIs were averaged together, as there are no differences in surface or performance between the left and right horizontal meridian (Himmelberg et al., 2023b). We conducted (two-tailed) Pearson correlations to evaluate the relation between V1 surface area and presaccadic enhancement for horizontal, upward, and downward saccades at the individual observer level.

## Results

For each test location (*horizontal, lower, upper*) and condition (*fixation, saccade*), we fit performance (d’) as a function of target contrast with a Naka-Rushton function (Naka and Rushton, 1966) (*Methods – Quantification & Statistical analysis*). Contrast responses during fixation were highly comparable across test locations (**Figure 2a-c**; gray), also indicated by the absence of a difference between their asymptotic performance level d_max_ (*F*(2,14)=1.09, *p*=0.355). We conducted repeated-measures ANOVAs with the factors test location and condition. Consistent with a recent study (Li et al., 2021b), presaccadic attention did not affect the semisaturation constant C_50_ (*F*(1,6)=1.81; *p*=0.221), but significantly modulated the asymptotic performance level d_max_ (*F*(1,6)=29.63, *p*=0.001), consistent with a response gain change. Importantly, this modulation interacted with saccade direction (*F*(2,14)=15.63, *p*<0.001). Bonferroni-corrected post-hoc comparisons showed that saccade preparation enhanced d_max_ at the saccade target (relative to fixation) before horizontal (*p*<0.001; **Figure 2a**) and downward (*p*<0.001; **Figure 2b**) saccades. Crucially, there was no presaccadic response gain benefit before upward saccades (*p*=0.217; **Figure 2c**). Correspondingly, the presaccadic benefit Δd_max_ (d_max_ *saccade* - d_max_ *fixation*) significantly varied with saccade direction (*F*(2,14)=15.63; *p*<0.001). Post-hoc comparisons demonstrated no statistical difference between the benefit caused by horizontal and downward saccades (*p*=1.00; **Figure 2d**), whereas the benefit before upward saccades (when present) was significantly reduced compared to both horizontal (*p*=0.003; **Figure 2e**) and downward (*p*=0.004; **Figure 2f**) saccades. Note that this differential effect was present for all observers.

**Figure 2.**
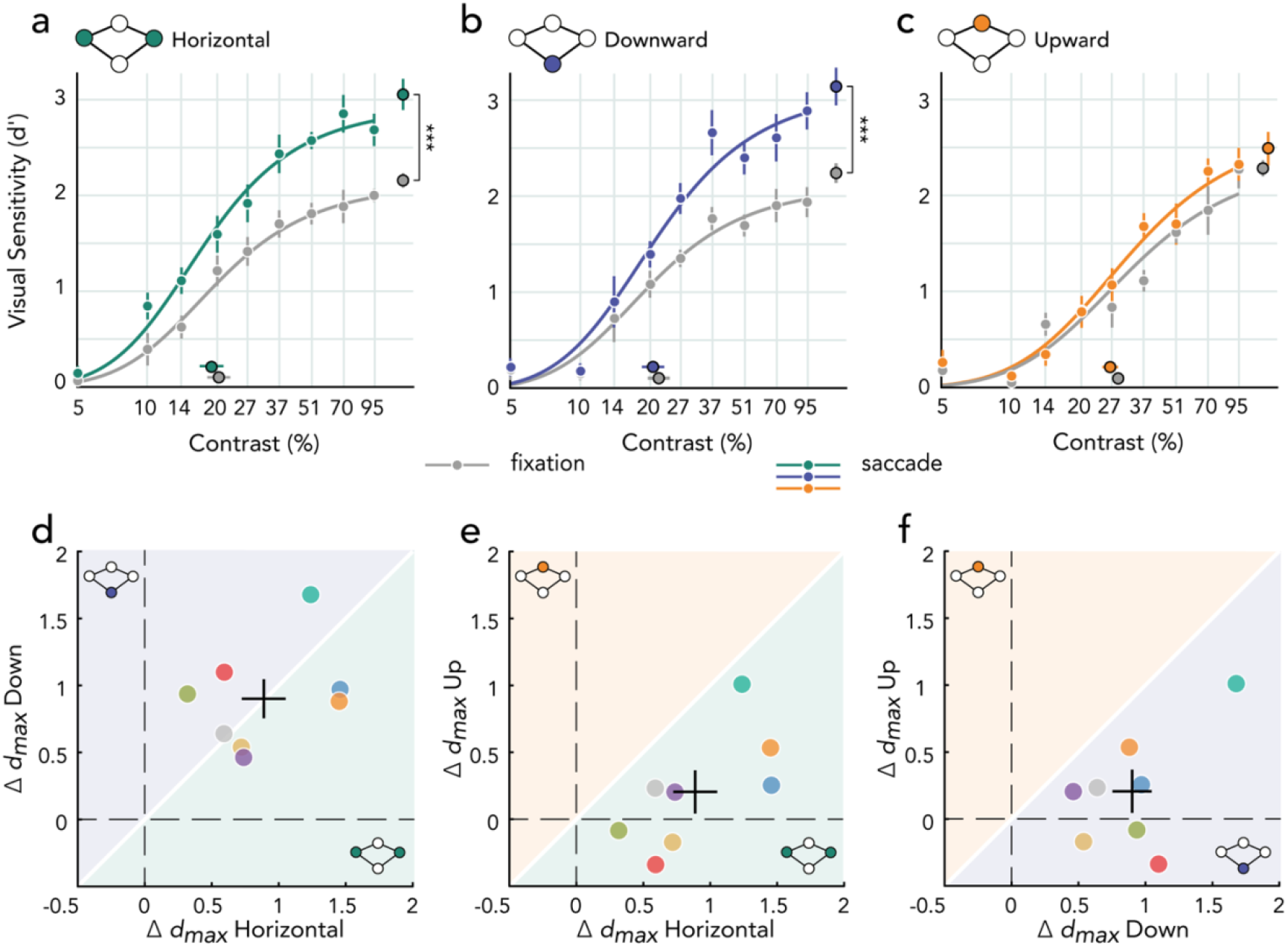
Contrast responses depending on saccade direction. Group-averaged psychometric contrast response functions (CRF; d′ versus contrast) measured during fixation (gray) or saccade preparation (colored) at the horizontal (**a**), lower vertical (**b**), and upper vertical (**c**) meridian. Group-averaged d_max_ and C_50_ extracted from individual observers’ CRFs are plotted at the right and the bottom of the figure. Error bars depict ±1 SEM. \*\*\**p*<0.001; (**d-f**) Pairwise comparison of individual observers’ presaccadic benefit Δd_max_ (d_max_ *saccade* - d_max_ *fixation*) at the horizontal (green background), lower (purple), and upper (orange) test locations. Black crosses depict the group average ±1 SEM.

We evaluated eye movement parameters to ensure that the missing presaccadic benefit before upwards saccades cannot be explained by differences in saccade latency or precision, e.g., comparatively slower or less precise upward saccade execution. A 2-way repeated-measures ANOVA with factors saccade direction and test contrast level showed a significant main effect of saccade direction (*F*(3,21)=8.94, *p*=0.006) on latency (**Figure 3a**), but no effect of stimulus contrast (*F*(8,56)=1.50, *p*=0.248) or an interaction effect between the two (*F*(24,168)=2.15, *p*=0.116). Post-hoc comparisons indicated significantly longer latencies for downward saccades compared to left (*p*=0.006), right (*p*<0.001), and upward (*p*=0.037) saccades. As this pattern was expected (Honda and Findlay, 1992; Tzelepi et al., 2010; Grujic et al., 2018; Hanning et al., 2022a; Kwak et al., 2023; Liu et al., 2023), we adjusted the delay between cue and test presentation to each observer’s horizontal and vertical saccade latencies (*Methods – Stimulus timing*) so that test presentation time relative to saccade onset was matched across saccade directions (**Figure 3d**); pairwise comparisons did not show any significant difference between upward saccades and those to other directions (all *p*≥0.106). Note that presaccadic perceptual enhancement was not correlated with saccade latencies for any saccade direction (all *p*≥0.671). Likewise, a saccade latency median split (within each participant) showed no significant difference in the presaccadic effect between relatively shorter and longer latencies for each saccade direction (all p≥0.467).

**Figure 3.**
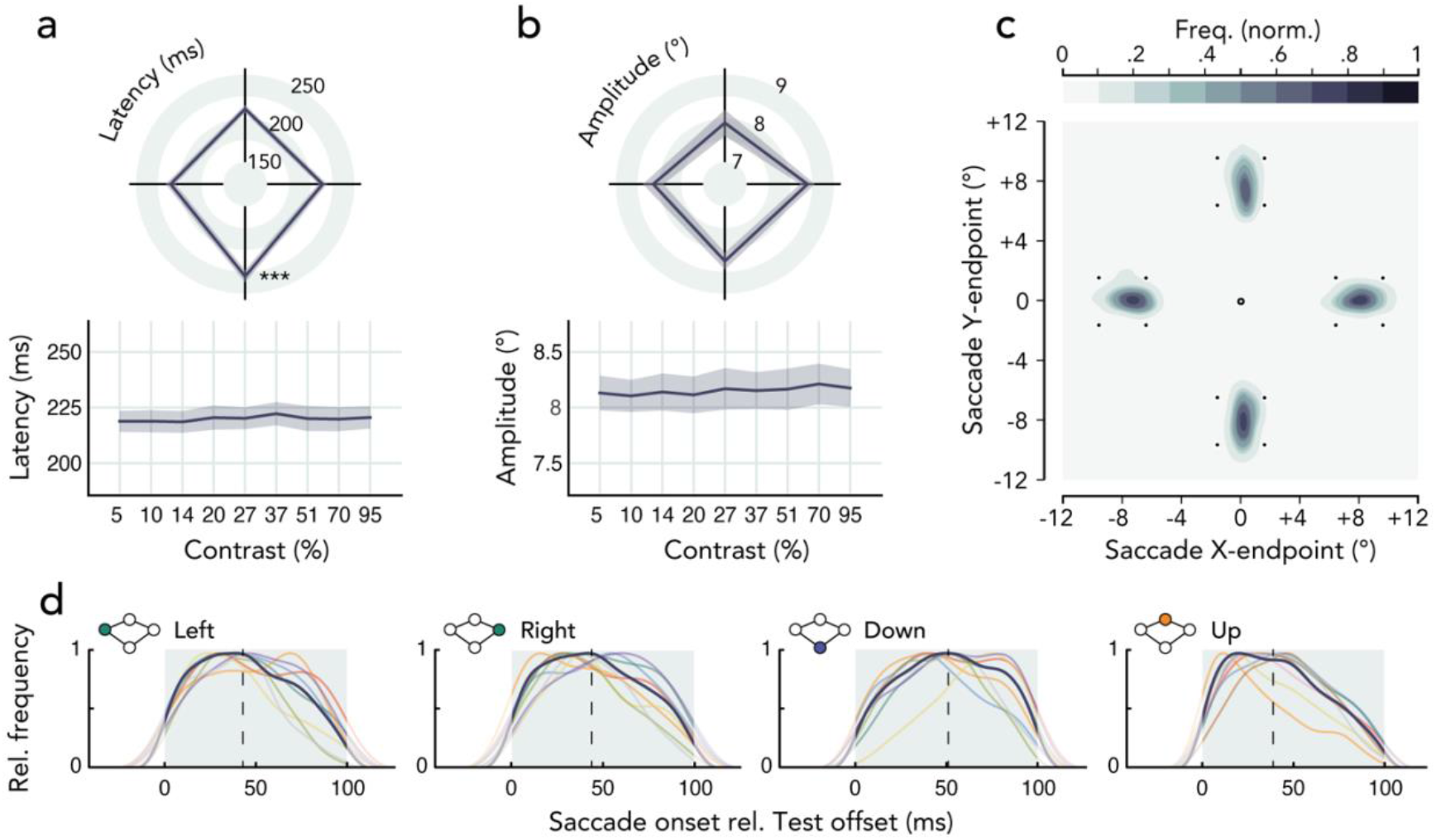
Eye movement parameters. Group average saccade latency (**a**) and amplitude (**b**) as a function of saccade direction (top) and test contrast level (bottom). Shaded error areas indicate ±1 SEM. (**c**) Normalized saccade endpoint frequency maps averaged across observers depicting saccade landing variance. (**d**) Density plots of saccade onset times relative to test offset, for each observer (thin colored lines) and across observers (bold line). Dashed vertical lines indicate the average saccade onset across observers; the gray background indicates the time window of trials included in the analysis (test offset ≤100ms before saccade onset).

Neither saccade amplitudes (**Figure 3b**) nor saccade landing errors (i.e., the mean Euclidean distance between saccade endpoints and saccade target center; **Figure 3c**) were affected by saccade direction (*amplitude*: *F*(3,21)=1.585, *p*=0.241; *landing error*: *F*(3,21)=1.292, *p*=0.306), stimulus contrast (*amplitude*: *F*(8,56)<1; *landing error*: *F*(8,56)=1.41, *p*=0.259), or their interaction (*amplitude*: *F*(24,168)<1; *landing error*: *F*(24,168)=1.37, *p*=0.265). In sum, latency and movement precision of upward saccades were comparable to those of horizontal and downward saccades, ruling out these parameters as source for the absent presaccadic benefit before upward saccades. In any case, the presaccadic shift of attention is not linked to the saccade landing position but to the *intended* saccade target (Deubel and Schneider, 1996; Stigchel and Vries, 2015; Wollenberg et al., 2018; Hanning et al., 2019b; Li et al., 2021b).

Our data show that presaccadic attention not only varies around the visual field, but also considerably among individual observers. Whereas observers consistently showed a presaccadic benefit of similar magnitude before horizontal and downward saccades (**Figure 4a**), we observed a mixed pattern for upward saccades, ranging from moderate presaccadic benefits to even presaccadic costs (i.e., lower presaccadic sensitivity at the upward saccade target than during fixation; 37.5% of observers). Note that for those observers displaying a presaccadic benefit at all locations, the benefit before upward saccades was lower than before horizontal and downward saccades.

**Figure 4.**
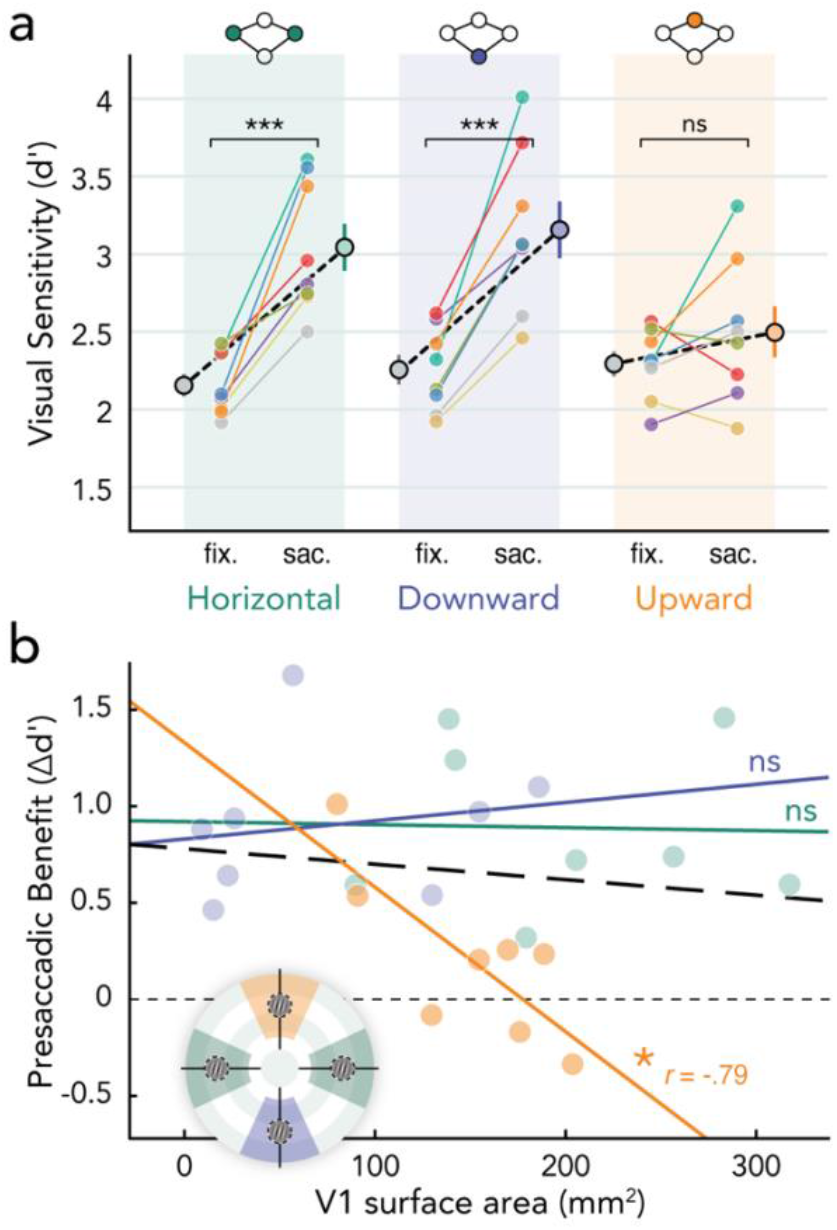
Linking presaccadic attention to individual differences in the surface area of primary visual cortex. (**a**) Individual observers’ d_max_ (extracted from their fitted CRF) for each test location and condition. Dashed black lines depict the group average, error bars depict ±1 SEM. (**b**) Individual observers’ presaccadic benefit (d_max_ saccade - d_max_ fixation) before horizontal, downward, and upward saccades plotted as a function of their respective localized measurements of V1 surface area. The circular icon depicts the extracted area: 4°-12° eccentricity, ±25° of radial angle extending either side of each cardinal meridian (note that horizontal surface area was computed as the average of left and right). \**p*<0.05. See **Supplemental Figure S1** for correlations between local V1 surface area and contrast sensitivity during fixation.

Notably, the visual field is non-uniformly represented in primary visual cortex (V1) –with greater surface area representing the horizontal than vertical meridian, and lower than upper vertical meridian (Silva et al., 2018; Benson et al., 2021; Himmelberg et al., 2021, 2022, 2023a). This directly reflects (Himmelberg et al., 2020), and likely (at least partially) underlies (Himmelberg et al., 2022), perceptual differences around the visual field (**Figure 1a**). Individual differences in contrast sensitivity at perceptual threshold (measured during fixation) correlate with the amount of V1 surface dedicated to processing the respective region of visual space (Himmelberg et al., 2022). Thus, higher contrast sensitivity at the horizontal meridian is a perceptual consequence of more dedicated V1 surface area. We confirmed that this link also holds in our study by correlating contrast sensitivity measurements obtained from 6 of the 8 observers at perceptual threshold in our previous study (Hanning et al., 2022a) with their V1 surface area measurements (using the same parameters as Himmelberg et al., 2022; see **Supplemental Figure S1**). We replicated the positive correlation between V1 surface area and contrast sensitivity during fixation (Himmelberg et al., 2022; Jigo et al., 2023), which provides further support for the notion that targeted fMRI measurements are robust even with small sample sizes (Himmelberg et al., 2021). Could this relation between primary visual cortex and contrast sensitivity also hold for presaccadic perceptual modulations? To explore whether presaccadic attention benefits are linked to V1 surface area, we correlated each observer’s presaccadic benefit before horizontal, downward, and upward saccades with their amount of V1 surface area representing the respective saccade target region. Individual observers’ surface estimates were obtained in a separate fMRI experiment (Himmelberg et al., 2021) (*Methods – fMRI Retinotopy & Surface computation*).

For horizontal and downward saccades, which showed the typical presaccadic sensitivity enhancement, we observed no correlation between presaccadic benefits and the amount of V1 surface area representing the saccade target (*horizontal* target: *r*(6)=-0.03, *p*=0.947; *downward* target: *r*(6)=-0.17, *p*=0.681). In contrast, inter-individual variations in the presaccadic effect preceding upward saccades were negatively correlated with the amount of V1 surface dedicated to processing the upward saccade target (**Figure 4b**). Thus, for individual observers, the less cortical surface representing the saccade target, the larger the presaccadic attentional benefit (*upward* target: *r*(6)=-0.79, *p*=0.019). This finding documents a link between cortical anatomy and the magnitude of presaccadic perceptual modulations (or lack thereof) preceding upward saccades at the individual observer level.

To underscore the significance of this finding, we have validated the consistency of the observed inverse relation between V1 surface area and perceptual presaccadic enhancement at the upper vertical meridian by evaluating data from our previous psychophysical study measuring presaccadic contrast sensitivity (Hanning et al., 2022a). Observers for whom fMRI retinotopy data was available from the NYU Retinotopy Dataset (n=5 independent, new observers) show the same pattern of a negative relation between V1 surface area and presaccadic benefit at the upper vertical meridian: **Figure 5** shows the pre-saccadic enhancement, now computed as the ratio of pre-saccadic contrast sensitivity, indexed at d_max_ (present study, n=8) or contrast threshold (previous study (Hanning et al., 2022a), n=5) and the respective measurement during fixation. Combined, these data from 13 observers confirm the inverse relation between V1 surface area representing the upward saccade target and the corresponding (diminished) pre-saccadic enhancement (*r*(11) = -0.71; *p* = 0.007).

**Figure 5.**
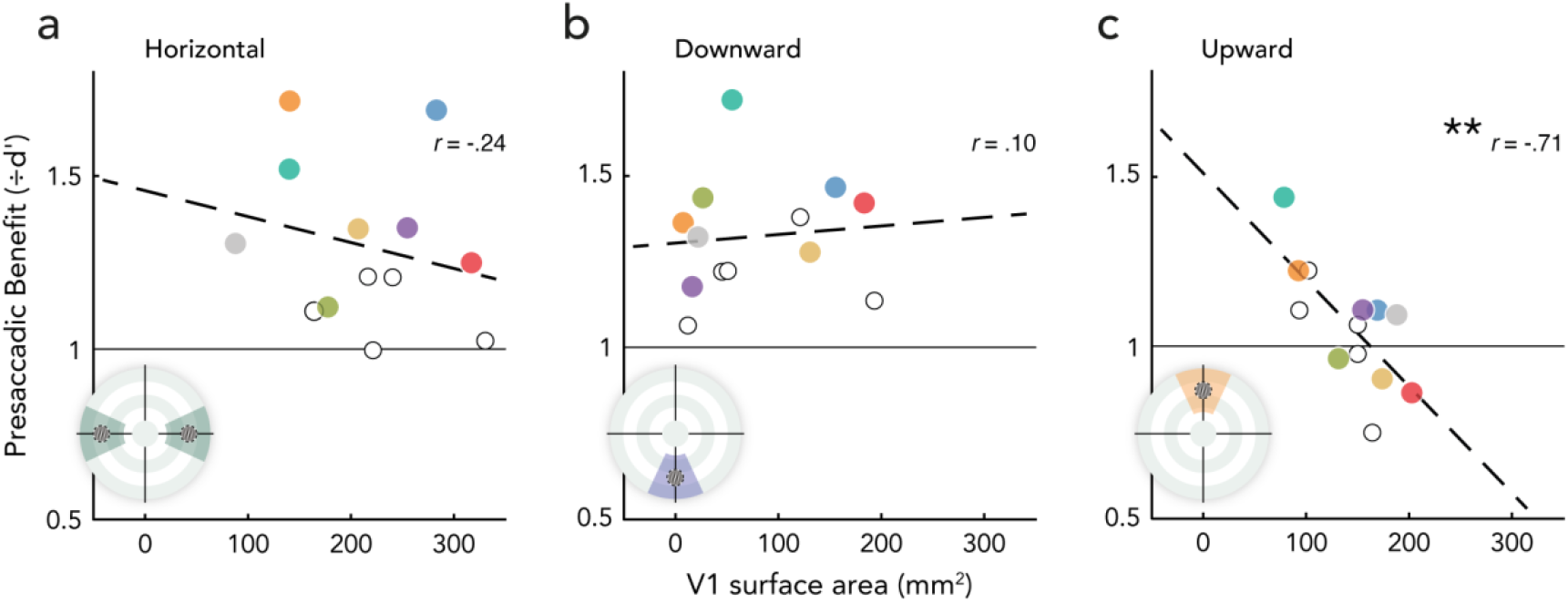
Presaccadic benefit in asymptotic and threshold contrast sensitivity (CS) as a function of local V1 surface area representing the respective saccade target region. Benefit computed as the ratio of asymptotic CS d_max_ saccade / CS d_max_ fixation (colored dots, matching n=8 observers in Figures 2-4) or CS threshold saccade / CS threshold fixation (white dots, n=5). Circular icons depict the extracted area: 4°-12° eccentricity, ±25° of radial angle extending either side of each cardinal meridian (note that horizontal surface area was computed as the average of left and right). \*\**p*<0.01.

## Discussion

To investigate how presaccadic attention for different saccade directions impacts the contrast response function we have optimized the detection of presaccadic benefits as a function of saccade direction: We (I) measured presaccadic contrast sensitivity across the full contrast range to capture the response gain effect by which presaccadic attention modulates contrast sensitivity; (II) adjusted overall task difficulty during fixation –which served as a baseline condition to quantity the effect of presaccadic attention– to compensate for performance field asymmetries; and (III) adjusted the test presentation time separately to each observer’s direction-specific saccade latency to ensure a measurement right before saccade onset, when presaccadic attention has its maximum effect.

We found that horizontal and downward saccades increased response gain, consistent with the well-established sensitivity benefit at the saccade target. Before upward saccades, however, presaccadic benefits were absent at the group level –and inconsistent across observers–, resembling our findings at perceptual threshold (Hanning et al., 2022a). We verified that differences in saccade latency, amplitude or landing precision (between observers or saccade directions) could not explain the diminished to absent upward presaccadic enhancement; some observers even showed a cost. Interestingly, we identified a neural correlate for this differential effect: Observers with a comparatively greater upward presaccadic benefit have relatively less V1 surface area dedicated to processing that target region.

The absent presaccadic benefit before upward saccades contradicts the notion that presaccadic attention is automatically deployed before any saccade; however, presaccadic attention studies typically rely on measurements along the horizontal meridian or do not evaluate potential differences among tested saccade directions (review: Li et al., 2021a). Likewise, neurophysiological measurements of presaccadic attention have not been evaluated as a function of saccade direction. Our present and recent (Hanning et al., 2022a; Liu et al., 2023) findings demonstrate that observations made for one saccade angle do not generalize to other directions and call for an investigation of presaccadic attention for other properties and tasks as a function of polar angle.

Although the strong coupling between saccade planning and visual attention has been undisputed, the *direction* of this relation is still debated. Proponents of the *premotor theory of attention* argue that visual attention is a product of the motor system; even covert shifts of attention, during fixation, result from (eye) movement planning (Rizzolatti et al., 1987; Craighero et al., 1999) (review (Smith and Schenk, 2012)). Oculomotor brain structures (e.g., FEF, SC) are also selectively modulated during covert attention tasks in human- and non-human primates (Nobre et al., 2000; Bogadhi et al., 2018; Bollimunta et al., 2018). However, distinct neuronal populations within these regions underlie covert attention and saccade preparation activity (Ignashchenkova et al., 2003; Müller et al., 2005; Gregoriou et al., 2012; Messinger et al., 2021). In sum, recent research refutes the view that the control of spatial attention is dependent on oculomotor control circuits (Hanning et al., 2019b, 2023; Hanning and Deubel, 2020; Masson et al., 2020; Messinger et al., 2021; review: Li et al., 2021a) and suggests the reverse: Goal-directed movements depend on preceding attentional selection to specify motor target coordinates – *selection-for-action* (Schneider, 1995; Deubel and Schneider, 1996; Baldauf and Deubel, 2010). Our results are inconsistent with both accounts: (1) Eye movement parameters for upward saccades are no different from horizontal and downward saccades; given that upward saccade motor programming is “typical” (has comparable saccade latency and precision), according to the premotor theory of attention it should cause a regular shift of attention to the motor target; (2) According to the selection-for-action account, the presaccadic shift of attention functions as a motor target marker and is a prerequisite for saccade execution. Here we show regularly executed upward saccades without a measurable attentional marker of motor target selection.

Assuming presaccadic attention is required for motor target selection but absent prior to upward saccades, how can we perform (accurate) upward saccades? Could presaccadic attention be deployed to upward saccade targets, without resulting in the typical corresponding perceptual advantages? This possibility could be tested by measuring neurophysiological correlates of presaccadic attention. Event-related potentials (ERPs), which track covert attentional (Luck et al., 1994; Luck and Yard, 1995) and motor target selection (Baldauf and Deubel, 2009) via sensory-evoked P1/N1 components, could reveal a dissociation between perceptual and neurophysiological correlates of presaccadic attention at the upper vertical meridian.

This is a plausible scenario given that sensitivity at the upper vertical meridian is lower than at other isoeccentric locations due to anatomical constraints in retina and visual cortex (Himmelberg et al., 2023b). Perceptual polar angle asymmetries are likely explained by the distribution of V1 tissue (Benson et al., 2021; Himmelberg et al., 2022, 2023a; Kupers et al., 2022), which parallels perceptual sensitivity differences around the visual field (Benson et al., 2021), even at individual observer level (Himmelberg et al., 2022). Could the reduced neural resources devoted to processing the upper visual field (partially) explain why saccade preparation fails to improve sensitivity at the upper vertical meridian? The absent group-level benefit at the upper vertical meridian could be related to its smallest surface area representation, largest receptive fields, and lowest preferred spatial frequency (Aghajari et al., 2020; Broderick et al., 2022). Our 4-cpd test stimulus may be less suited for driving neural responses at the upper vertical than the horizontal meridian (Jigo et al., 2023) at the 8° eccentric saccade target.

At the individual level, however, observers with comparatively less V1 surface representing the upward saccade target had a relatively greater –but compared to other directions still severely reduced– presaccadic benefit. According to the above explanation, however, one would expect observers with relatively larger surface area to have larger presaccadic benefits. Presaccadic attention causes the neural representation of a target stimulus to become more ‘fovea-like’ and likely sharpens receptive field sizes at the target location (Li et al., 2016, 2019; Ohl et al., 2017; Kroell and Rolfs, 2021; Kwak et al., 2023). If observers with less surface area representing the upper vertical meridian have larger receptive fields, and presaccadic attention sharpens them, this could result in a (relatively) greater presaccadic benefit for observers with smaller cortical upper-vertical visual field representation.

This effect was not observed at the other cardinal locations; computations of neurons encoding the upper vertical meridian may differ from those at the other meridians. Magnifying the stimulus size according to cortical representation eliminates contrast sensitivity differences across eccentricity but not polar angle (Jigo et al., 2023), suggesting that polar angle performance differences are likely mediated by neurons with differential image-processing capabilities (i.e., differently tuned spatial filters). Using reverse correlation, we have shown that higher sensitivity to task-relevant orientation and spatial frequency and lower internal noise at fovea than perifovea underlie eccentricity-dependent variations (Xue et al., 2023). Polar angle differences, however, seem to stem from different orientation and spatial frequency tuning functions; e.g., the upper vertical meridian is less sensitive to orientation and tuned to low spatial frequencies (Xue and Carrasco, 2023).

Our results challenge the mandatory link between saccade programming and attention, and raise the question how a smooth transition from (presaccadic) blurry peripheral information to its high-acuity equivalent (once foveated after the saccade) is achieved also across upward saccades: Transsaccadic integration is assumed to rely on predictive presaccadic sensitivity modulations to render peripheral representations at the saccade target fovea-like (Herwig, 2015; Rolfs, 2015). Interestingly, contrast sensitivity is reduced at the postsaccadic center of gaze *after* upward saccades (Liu et al., 2023). This effect could result from a diminished preparatory shift of presaccadic attention to the upward saccade target. Likewise, perisaccadic perceptual mislocalization persists longer after upward saccades than saccades in other directions (Grujic et al., 2018) – which has been linked to asymmetries in SC neural properties representing the upper vs. lower visual hemifield (Hafed and Chen, 2016). This study, however, does not record from neurons with receptive fields at the upper vertical meridian (±30°), where our analyses focus. Future work should document whether reported asymmetries also extend to the vertical meridian, where perceptual and cortical asymmetries are more pronounced (Himmelberg et al., 2023b).

We have recently investigated presaccadic acuity thresholds, which were equally increased (compared to fixation) for all cardinal saccade directions, including upward saccades (Kwak et al., 2023). This saccade-direction unspecific effect is seemingly inconsistent with the direction-specific effect of presaccadic attention on contrast sensitivity. However, acuity and contrast sensitivity measurements targeting different points on the contrast sensitivity function (CSF), which is characterized by a joint manipulation of contrast and spatial frequency. Whereas visual acuity, measured at full contrast, corresponds to the cut-off point of the CSF, contrast sensitivity is measured closer to the peak of the CSF. If presaccadic attention reshapes the refhape ces CSF differently depending on saccade direction, this could explain its differential effect on (enhanced) acuity and (unaffected) contrast sensitivity at the upper vertical meridian.

To conclude, even when accounting for known polar angle differences in peripheral vision during fixation and testing in a contrast range optimized for detecting presaccadic sensitivity modulations, saccades made to the upper vertical meridian lack the typical presaccadic benefit. This effect may be related to individual differences in V1 cortical surface shown here, and to different computations underlying perceptual modulations at the upper vertical meridian. Our findings call into question the generalizability of behavioral and neurophysiological measurements obtained from one saccade angle to another. Future studies should test whether oculomotor feedback signals (between eye movement and visual areas), which are thought to underlie presaccadic modulations of visual perception, are different for upward saccades than those in other directions, and whether further behavioral correlates of presaccadic attention –such as orientation tuning or the selective shift of sensitivity to higher spatial frequencies– are diminished, or even absent prior to upward saccades.

## Author contributions

Conceptualization and methodology: NMH, MC; Software: NMH; Investigation: NMH; Overall analysis and visualization: NMH; fMRI analysis: MMH; Writing—original draft: NMH; Writing—review & editing: NMH, MMH, MC; Funding acquisition: NMH, MC.

## Data availability

Raw eye tracking and behavioral data will be available from the OSF database at https://osf.io/xrtbf/; preprocessed fMRI data is available from the openNeuro platform at https://openneuro.org/datasets/ds003787/versions/1.0.0;

## Conflict of interest statement

The authors declare no competing financial interests.

## Acknowledgements

This research was supported by a Marie Skłodowska-Curie individual fellowship by the European Commission (898520) to NMH and NIH NEI Grants R01-EY-019693 and R01-EY-027401 to MC. We thank the members of the Carrasco Lab, as well as Kate Salamatov and Monty Cox for helpful discussions.

## Supplemental Information

**Supplemental Figure S1.**
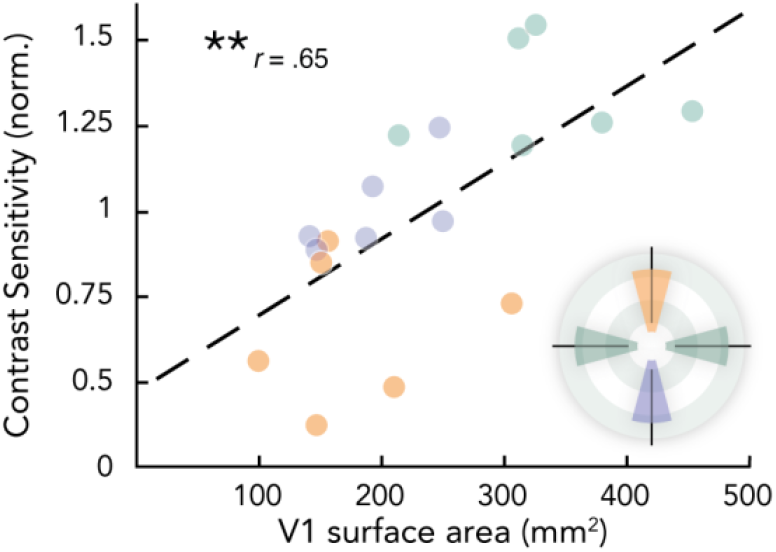
Local V1 surface area predicts contrast sensitivity during fixation. Individual observers’ normalized contrast sensitivity measured during fixation in the context of our previous study (Hanning et al., 2022a) as a function of the respective localized V1 surface area measurement. The circular icon depicts the extracted area: 1°-8° eccentricity, ±15° of radial angle extending either side of each cardinal meridian, matching the original study (Himmelberg et al., 2022). Note that horizontal surface area was computed as the average of left and right. \*\**p*<0.01.

**Supplemental Movie 1.**
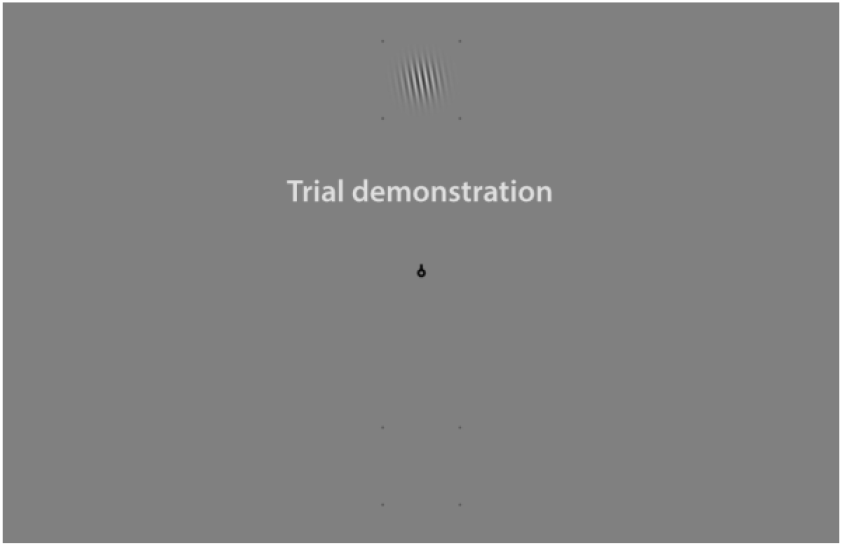
Video demonstration of an exemplary vertical *saccade* trial (grating contrast 70%).

## Notes

### Competing Interest Statement

The authors have declared no competing interest.

### Summary of Updates

Additional analyses visualized in Figure 5 and Figure S1 (for details see main text).

